# Cortical Spikes use Analog Sparse Coding

**DOI:** 10.1101/2020.10.19.331389

**Authors:** Dana H. Ballard, Ruohan Zhang

## Abstract

Quantifying the message communicated by neurons in the cortex by averaging action potentials over repeated trials of a given stimulus can reveal neuronal tuning features. For example, simple cells in the visual cortex have been characterized by reverse correlation based on the detailed structure of their oriented receptive fields. This structure, in turn, has been modeled using large libraries of such receptive fields to allow the simultaneous coding of visual stimuli with small numbers of appropriate combinations of cells selected from the library. This strategy, known as *sparse coding*, has been shown to produce excellent approximations for natural visual inputs. In concert with this mathematical development has been the discovery of cells’ use of oscillations in the gamma frequency range for general coding tasks, such as a mechanism for synchronizing distal networks of neurons. More recently, spikes timed with oscillations have been shown to exhibit local phase delays within a single gamma cycle, but such delays have resisted a behavioral functional interpretation. We show here that a specific coordinate system for the gamma cycle allows resultant phase delays to be interpreted quantitatively in classical terms. Specifically, extracted phase delays from mice viewing oriented sinusoidal grating images are shown to have the same distributions as those from a computer sparse coding model using natural images, suggesting for the first time a direct link between experimentally measured phase delays and model receptive fields.

**Significance Statement:** Networks of pyramidal cells in the cortex exhibit action potentials (spikes) that are characterized by randomness and low firing rates. Spike averaging methods have been ordinarily useful in dealing with these features to reveal behavioral task structure, but the randomness and slowness so far prevented the specification of a satisfactory generative spike model. We show that a spike can be analyzed using the context of a specific phase of the gamma component of its membrane potential. The result is each spike can be can be assigned a scalar, which makes it immediately useful for network models.

## 1 Introduction

The discovery that synchronous inputs could be efficacious in generating an action potential opened up new characterization of cortical circuits [Abeles, 1991], both in spatially organized networks [Singer and Gray, 1995, Engel et al., 2001] and the propagation of temporally synchronized signals [Ikegaya et al., 2004, Hayon et al., 2005]. Although such research focused on the role of a spike, subsequent experiments evidence showed that an action potential’s temporal reference may exploit the gamma frequency band (approximately 30 ~ 80 Hz). Specifically, several experiments have revealed its use in communication between distal cortical locations, [Fries, 2015, Fries et al., 2001, Chalk et al., 2010] including its modulation with specific stimuli [Hermes et al., 2019] and a diversity of phase relations [Maris et al., 2016]. While the main focus of such research has been on global synchronization, our present study focuses on the use of a spike’s temporal reference based on information contained within *local oscillations*. This more fine-grained approach explains a spike’s local phase delay with respect to a single spike encompassing oscillation’s wavelength as a scalar instead of a single digit, thus vastly improving both the information rate and reducing the space needed. Local phase coding in the cortex was suggested over twenty years ago [Hopfield, 1995], but the development of in vivo whole-cell patch-clamping [Hamill et al., 1981] has given researchers unprecedented quantitative access to neuronal membrane potential dynamics in awake behaving animals [Gentet et al., 2010], revealing for the first time the presence of low amplitude gamma oscillations within the membrane potential. That observation and high-frequency recordings allowed us to extract high-precision local spike phase shifts automatically from intracellular data obtained in awake animals [Perrenoud et al., 2016, Zhang and Ballard, 2020].

This paper describes a novel way of processing a cell’s membrane potential that allows the message carried by spike phases to be automatically extracted. Moreover its scalar structure suggests a direct correspondence with scalar responses of abstract cell models. Specifically, we show that information in sets of delays is close to identical to the information in Sparse Coding models for simple cells in the striate cortex [Olshausen and Field, 1996b]. Sparse Coding is an attractive model choice for its elegant mathematical rationale. The model accounts the stimulus coding as trading off the accuracy of the receptive fields with the cost of the number of cells required by the model to perform its computation. The model also provides a close fit to the observed distributions of simple cell parameters such as size and spatial frequency [Olshausen and Field, 1996a].

For the local delay code, the paramount experimental analysis issue is if the local phase delay is to be measured, the action potential needs a time reference. We chose the phase statistics from patch-clamp data [Perrenoud et al., 2016], which consisted of membrane potential recordings from primary visual cortex of awake mice while observing images of moving gratings. Our software allowed us to extract the position of each action potential within its local gamma frequency profile. An overwhelming number of spikes occurred between the nearest preceding gamma minimum and the following maximum. We chose the minimum as a time reference and recorded the interval between the minimum and the position of the spike as an analog phase delay. The presupposition is that a cell’s synapses are experiencing a volley of such input delays, and the delay we measure is indicative of the cell’s resultant output. The assumption is that earlier spikes in the volley are more productive in generating a spike output with a short phase delay. We compared the local phase values from seven cells’ membrane potentials containing over 2,540 action potentials with an equivocally large number of model cell outputs. The comparison of the two distributions showed an extremely close match, suggesting a comparable functional interpretation for the phases in sparse coding of the receptive fields.

## 2 Results

Our model used previously-obtained intracellular recordings from seven patch-clamped pyramidal cells from a head-fixed awake mouse. These data were initially part of Perrenoud and colleagues’ earlier study [Perrenoud et al., 2016] and are freely available online on the Dryad repository. A session of the recording consists of a series of trials, each seven seconds long. Each trial starts with a two second-long stimulus-OFF baseline, followed by a 3-second presentation of a visual stimulus of a moving oriented sine grating, and finishing with a two second-long stimulus-OFF period. The task of holding on to a cell for many seven second trials can be delicate; in our database of seven cells, the number of trials per cell ranged from 39 trials up to 80. Each trial consisted of a specific moving grating. Eight orientations were used, and the orientation for each trial was chosen randomly. The number of action potentials per trial ranged from none to fourteen. Some of those spikes were outside of the 3-second interval. Our software extracted, all in all, 2,745 action potentials and their phases relative to the background gamma oscillations, but we analyze the 2,540 spikes extracted during the stimulus-ON period.

Although we could only measure one output spike at a time, each such spike is assumed to be part of a larger latent volley of input spikes from other cells used to code an output within that gamma cycle. However, even though experimentally measured synaptic input delays are not from the same volley within a single gamma cycle, we still use them. The volley properties can be obtained by sampling all the spikes irrespective of their generation times. This assumption is one of ergodicity where sampling in time is representative of sampling in space.

Next, we compared the distributions of phase delays with the distributions of sparse coding responses. The comparison assumes that the sparse coding model also uses parallel signal components. This decision is motivated by the theoretical constraint that the responses for each input image must be chosen probabilistically to conform to Bayesian principles [Doya, 2007, Jehee et al., 2006a]. Thus we combined the model projections as if they were generated in parallel. The sparse coding samples are comparable with the experimental samples since both are ergo chosen probabilistically.

Both distributions represent the coding of similar image information: oriented grating images responses in the experimental dataset, and image patches sample from natural outdoor images in the modeling dataset. Nevertheless, the comparison of the two distributions is derived from very different contexts. The model projections were computed using a parallel version of sparse coding, whereas the spike phase distributions were derived from gamma filtered membrane potentials. The two distributions shown in Figure 1a,b have very different venues that have different conventions, but very little processing needs to be done to compare them. Spike phase distributions were derived from their contextual locations of gamma filtered membrane potentials, and the shortest delay measured from the initial gamma minimum resulted in the largest response. Thus, with the progression on the time axis, the responses are made exponentially less as a function of delay length. In comparison, the model projections using parallel sparse coding were normalized so that the largest one was unity, and a frequency normalized of the projection is plotted with a normalized histogram representing the probability density function of the model data. Thus both plots represented the frequency of responses, oriented so that each of the largest responses are located at their origins.Next, each normalized histogram was used to create a cumulative probability density function.

**Figure 1:**
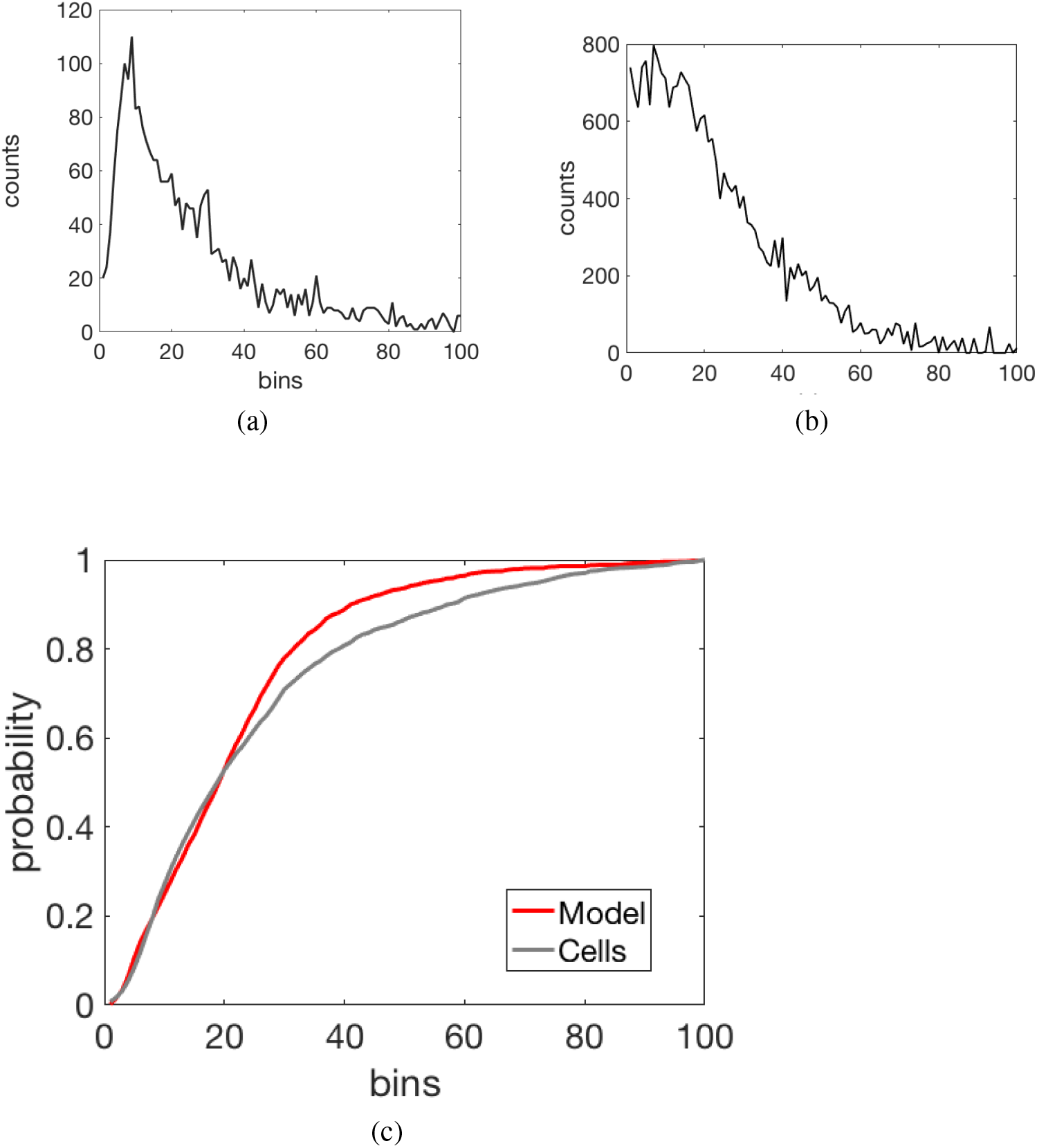
Comparing experimental data with model responses. (a) Experimental data: The responses generated by 2,540 action potentials, coded as phase delays relative to the gamma oscillations, are grouped into a histogram. If spikes occur probabilistically, then the cells with the larger input projections should be chosen more frequently and their delay should also be shorter. (b) The responses from the model coding many simulated image coding trials can be plotted as a histogram (c) The cumulative probability distributions for the experimental data (grey) and model(red) show a very close match.

The final step is to facilitate their comparison by comparing the two cumulative distributions. These two distributions were found to be very similar, as shown in Figure 1c.

A straightforward test is Kolmogorov-Smirnov [Massey Jr, 1951], which tries to reject the distributions as being dissimilar. The parameter measured is the maximum difference between the two curves. We used Matlab’s *kstest2*. The result is that the test produces a p-value of 0.2606, implying that there is a 74% probability that the two distributions of data and model can be thought of as similar. The conclusion is that the sparse codes representing image data distributions are nearly identical to the phase delays distributions representing the same image coding task.

That this match is so close is even more significant in the light of the variations seen in the filtered membrane potentials. Figure 2 shows that the gamma amplitudes vary by almost a magnitude and that the gamma frequencies cover the whole range of the low gamma band (33-58 Hz). The fact that the distribution of phase delays is invariant to these ranges argues in favor of using the phases as a coding signal. One might argue that this background variation would be introduced in the experimental distribution, but the statistical test argues that this is not the case.

**Figure 2:**
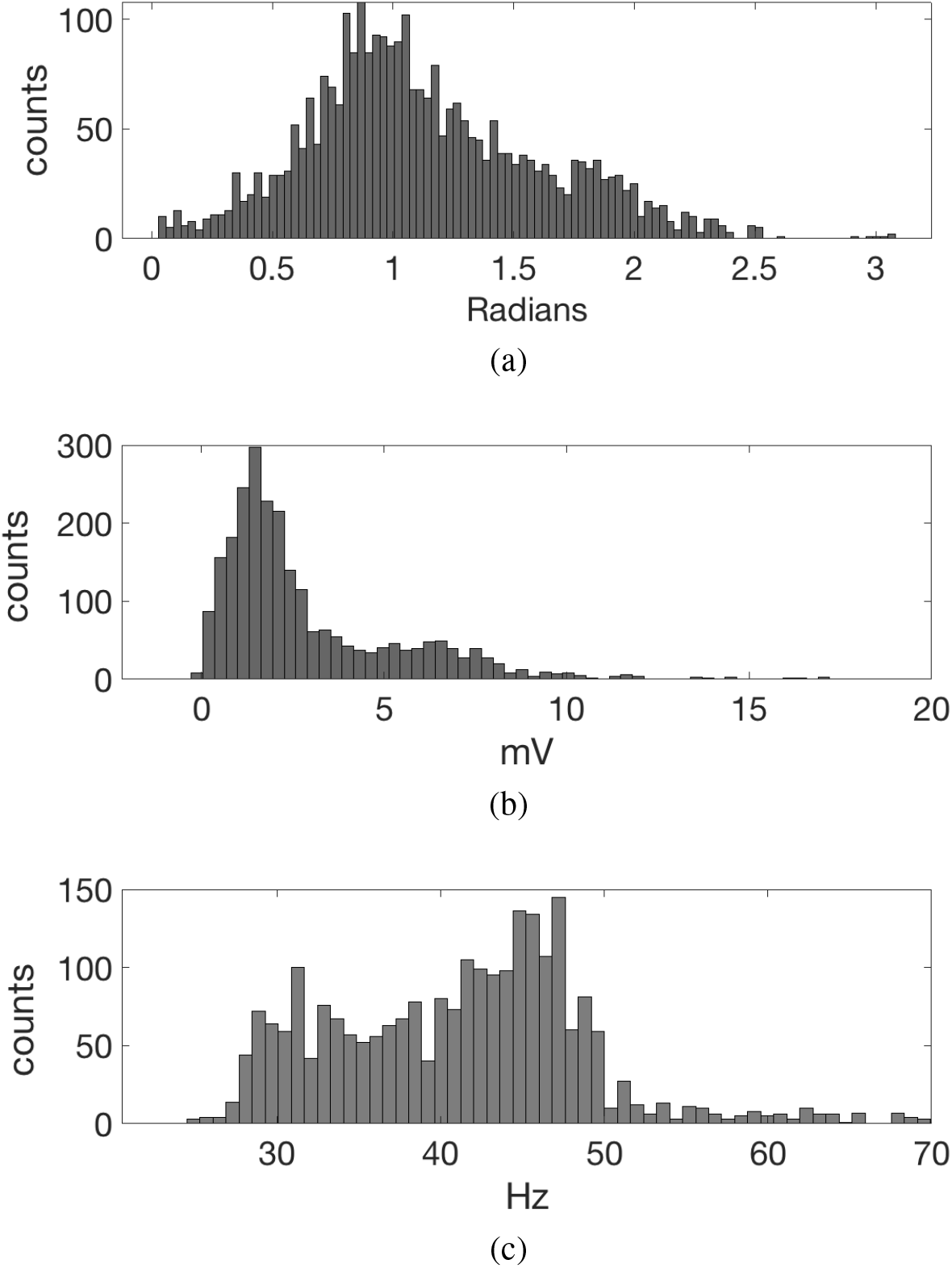
The phase delays extracted prove to be invariant to variations. (a) **delay coordinates** The great majority of the action potentials lie in the first half of the encompassing gamma cycle and most of those have a delay near one radian. Once these are exponentiated the resultant distribution matches the sparse coding model distribution closely, implying that the phases are invariant to the variations in gamma amplitude and modulating frequency. This data includes all seven seconds of each trial.(b) Distribution of the gamma amplitude at each action potential. Although the most common excursion between the filtered gamma amplitude is ±1 millivolt, the excursion can be as high as ± 9 millivolts. (c) Distribution of frequencies. The bulk of frequencies are evidenced in the low gamma range of 33 to 58 Hz

## Methods

The methods are grouped into separate experimental and model sections.

### Computation of the phase delay for a spike

Five steps were used to extract the local phase delays from the membrane potentials of seven cells.

1. The action potential components were removed from the the membrane potentials using the quadratic extrapolation method [Perrenoud et al., 2016];
2. The membrane potentials were bandpass filtered to extract the gamma frequencies in 33-58Hz range;
3. Each action potential was analyzed in the context of the filtered gamma membrane potentials to extract its associated phase delay;
4. Those delays were converted to a response (an analog quantity) using our model;
5. Their distributions were normalized to compute a cumulative probability distribution function.

The part of the membrane potential defining a spike must be removed before gamma filtering. If this is not done, the relatively large spike amplitude will unduly influence the gamma filtering result. The method we chose, that [Perrenoud et al., 2016] interpolates the membrane potential based on a smooth fit to the nearby potential either side of the spike. We chose ± 2*ms* as defining the interval straddling the spike to be interpolated as giving highest rates of phase-detection across the database of cells. Figure 3 shows the result of computing the portion of the action potential to be removed in red. The software to detect the related delays addresses each spike in turn.

**Figure 3:**
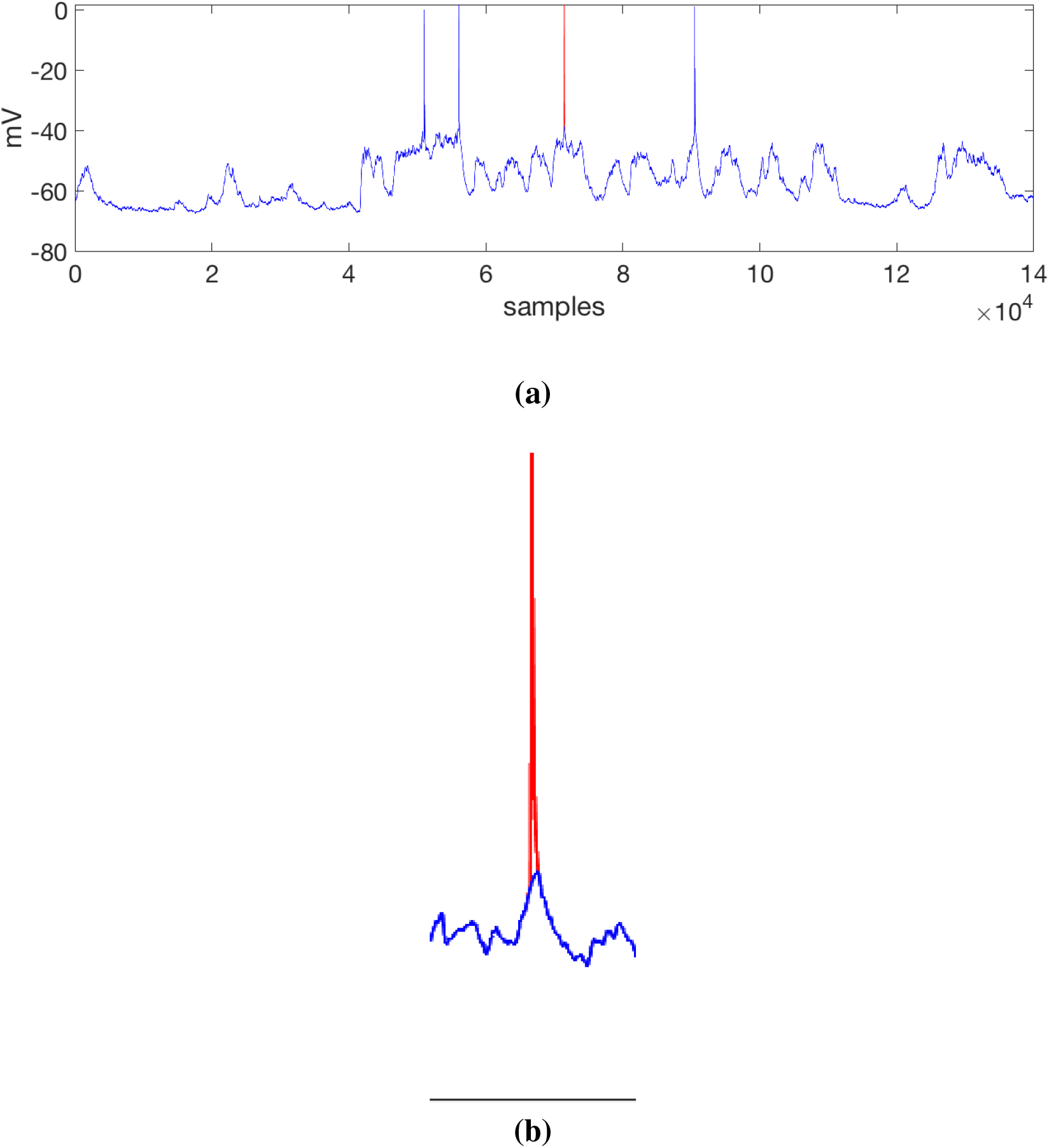
Spike removal. (a) Membrane potential for four action potentials when the third one showing the spike potential in red that will be moved. (b) An inset showing the interpolation detail. The width of the spike base used by the interpolating processes is 7.5 milliseconds.

Once the spike has been removed from its membrane potential vicinity, an area of a one-half second straddling the spike location is filtered with a Butterworth filter, whose parameters are shown in Table 1. The filter parameters are tuned to isolate the component of the signal in the low gamma range. Longer filter intervals make no difference in the gamma potential near the spike.

**Table 1:**
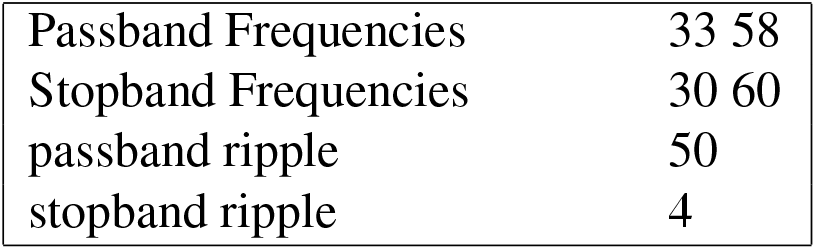
Butterworth filter parameters (in Hz) The input to the filter is one second of the membrane potential centered on an each action potential. The filer is designed to pass a low gamma band in the interval [33 58] Hz.

Figure 4 illustrates an example 7-second long whole-cell patch-clamp recording trial from a mouse V1 pyramidal cell. The membrane potential was filtered to isolate the gamma frequency band, and then for each action potential, we extracted its phase relative to the underlying gamma cycle. Each such phase delay was transformed into a response (an analog quantity) using an exponential decay function.

**Figure 4:**
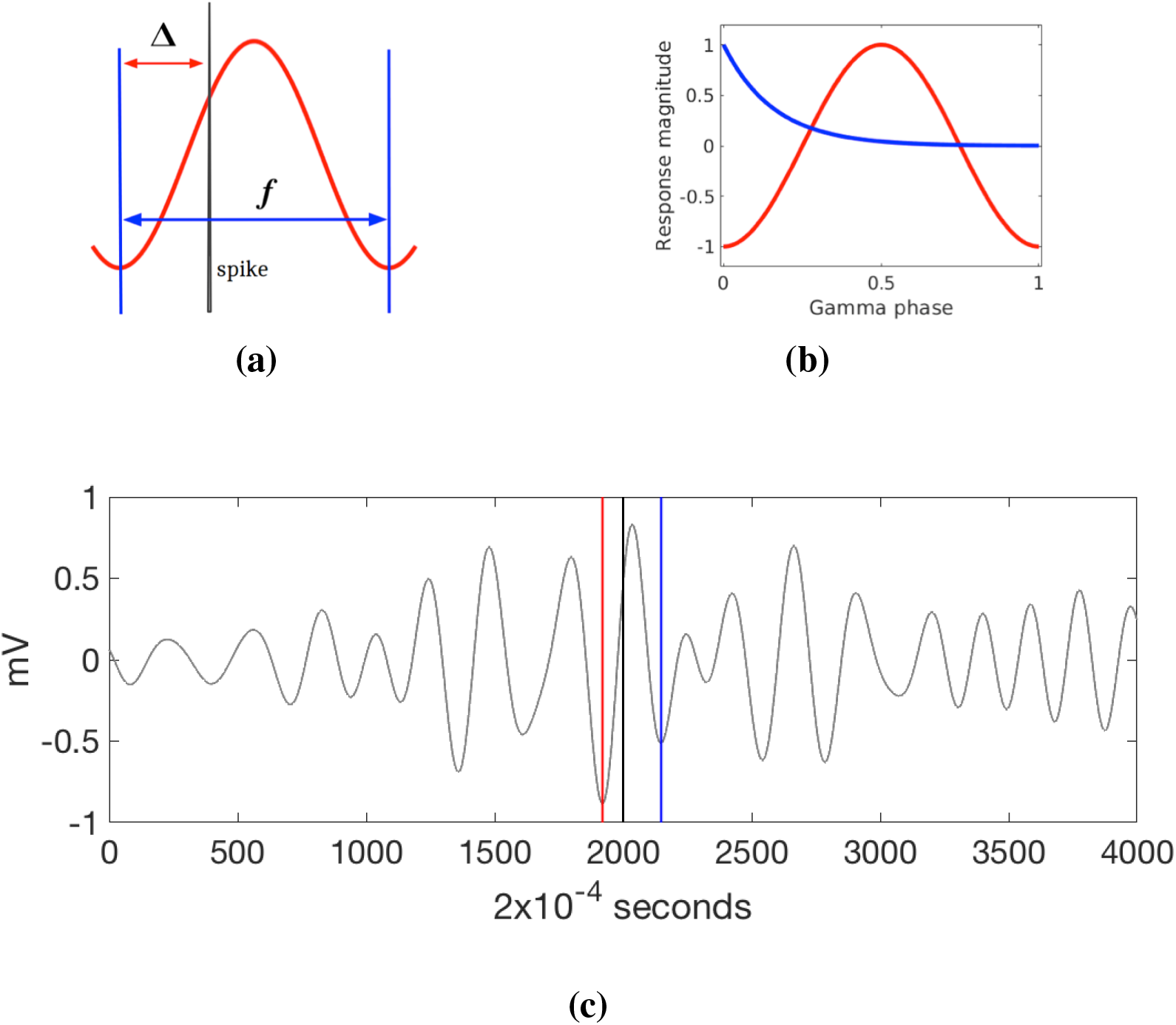
Information extracted from gamma modulation. Once membrane potential is frequency filtered in the gamma passband, it can be paired with the action potential to extract features related to its phase delay.((a) The spike location’s leading gamma local minimum becomes the zero for a local coordinate system. The time between this zero and the spike is the phase delay. Next, the minimum after the spike is also determined, allowing the local frequency to be estimated as the reciprocal of the wavelength distance between the two minima. This allows the delay to be measured in radians.(b) To interpret the value represented by the action potential, an exponential function is used. A spike signals an analog quantity (blue) coded as a phase delay from the trough. Short delays represent large magnitudes.(c) The process of extracting the phase delay features uses one half second gamma potential that straddles the action potential. Here the location of the leading minimum is shown as a red vertical. The action potential position is marked in black. The trailing minimum is marked in blue. The 20 samples per millisecond provide a good resolution for these measurements.

The gamma phase coding model requires choosing a time reference, i.e., the gamma cycle starting point. Selecting the first gamma minimum immediately preceding the spike as this point worked best, as almost all of the spikes fell within this half of the gamma cycle, and the delays were in the range of approximately 0 ~ 16 ms. Other important measures extracted were the wavelength, measured as the distance between successive minimums, as well as the maximum gamma amplitudes near the spikes.

These measures were extracted from every action potential. A typical experiment per cell can consist of 80 trial recordings that are each seven seconds in length and approximately 300 spikes in all. Our software automatically extracted for each spike, its interval, frequency, and location in the trace. Figure 5a shows the instantaneous frequencies for 80 trials of a single cell consisting of 264 spikes in total. For the same cell, the intervals for all the spikes are extracted and plotted with the same colour convention in Figure 5b.

**Figure 5:**
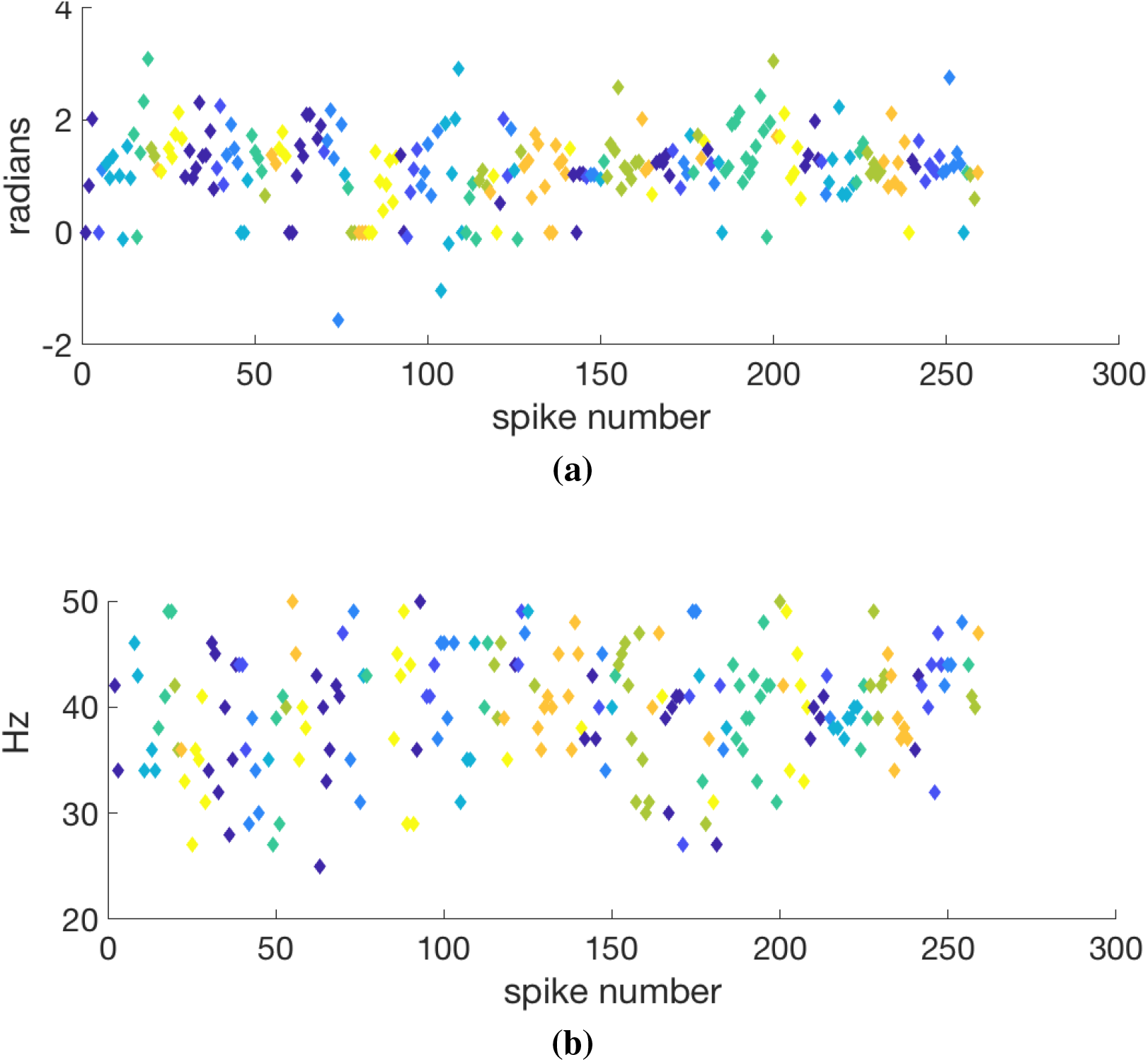
Somatic potential multi-trial analysis for a single cell with 80 trials. Each symbol denotes a specific spike. Each trial is associated with a particular oriented visual stimulus, and the spikes from that trial use the same color. A total of 260 spikes were detected. For each trial, the gamma filtering shown in Figure 4 is applied and the local gamma analyzed to extract its phase interval Δ and frequency *f*. (a) The spike data are plotted in the order of their occurrence. A few minimal negative phases occurred just before the chosen origin of a gamma minimum and are adjusted by moving the origin accordingly. The two points with large negative lead times are ignored as outliers. Fig. 2a shows the adjusted curve. (b) The extracted frequencies for the intervals are shown in the top figure. Some of the local wavelengths were challenging to extract to local variations in the waveform and are not shown.

#### Computing the experimental distribution

Each of the experimental examples’ coding cells has a response that conveys how well the current grating image input matches its receptive field. At any instant, we can only measure one spike phase delay, but since we assume from the sparse coding model that the participant cells in a volley need to be chosen probabilistically, we invoked an ergodic assumption to combine the phase data for any spike that had the same grating input. Thus we simulated the data from a volley using the phase delays from a large number of trials.

At the final step, the output phase values were converted to responses. This conversion is an open question. As discussed in the model, we used an exponential function where phase delay of zero measured at the gamma minimum was translated into a response of unity and a subsequent delay *d* was transformed into a response *r* by:

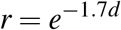

the factor of 1.7 was chosen to obtain the best fit to the model responses distribution. With this transformation, we are assuming that the distributions of experimental cells’ phase delays are commensurate with the distributions of model cells’ projections.

### Sparse coding volley model

Four steps were performed to obtain the sparse coding response distributions.

1. The natural images were filtered appropriately for the lateral geniculate nucleus;
2. The receptive fields were learned from a database trained on natural images using the algorithm of Lee et al. [Lee et al., 2007];
3. Fifty receptive fields were sampled probabilistically for each input image using our parallel algorithm;
4. Their distributions were scaled appropriately.

Since the experimental patch-clamp data uses oriented moving grating images for visual input, a good choice of model is one that can account for the properties of simple-cell oriented receptive fields. As previously introduced, the best choice for a pyramidal cell’s functionality is that of sparse coding, where a reduced set of cells chosen from a substantial surfeit can explain the parameter distribution of simple cells’ receptive fields [Olshausen and Field, 1996a]. It has been shown to generalize to cortical hierarchies via predictive coding [Rao and Ballard, 1999].

However, spikes generated in the context of gamma frequencies present us with a crucial problem. The classical sparse coding algorithm is sequential. Initially, a cell is chosen for an approximation for the input and subtracted from it, leaving the residual to be fit with another cell. This process of selecting cells from an evolving residual continues until a suitable approximation is achieved. However, this strategy is not practical with a gamma based code as computing and selecting each cell would need a gamma cycle and thus would be too slow to be viable.

To solve this problem, we developed a parallel sparse coding algorithm that generates the code in one parallel volley during one cycle of a specific gamma frequency. The requisite neurons in the volley must be selected probabilistically, each with odds proportional to their receptive field projections. Selecting Probabilistic selection is necessary to ensure that all neurons participate in the receptive field learning process appropriately.[Jehee et al., 2006b]

Given a target stimulus, a neuron’s response magnitude *r_i_* is calculated as a function of the input’s projection onto the synapses of the receptive field it represents. A standard choice of probability function is the Boltzmann distribution where a cell is chosen with probability

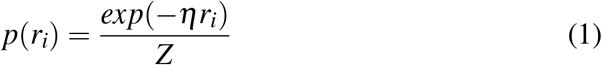

where *η* is a temperature parameter, and *Z* is a normalizing factor. The input’s projections’ relative magnitudes determine a neuron’s probabilities to be part of the representation. By exploiting lateral connections between neurons, the normalization can be done in one parallel step within a gamma frequency cycle. Our particular sparse coding implementation uses fifty coding cells for each input, chosen probabilistically from a base of one thousand candidates. The number of cells chosen represents a balance between being sparse, approximately 25% of the 196-pixel input, and providing a good representation [Zhang and Ballard, 2020]. A distribution of one thousand image samples, each with 50 sparse responses, was computed from this setting to compare with real cell data.

Our parallel algorithm is described pictorially in Figure 6. 14 × 14 lateral geniculate nucleus (LGN) images have been filtered with a transform appropriate for their transit from the retina. Then these images can be represented with 50 coding cells selected in parallel. Representing the image as a vector **I**, each potential coding cell as a vector **u**_*i*_ has an associated response *r_i_* = **I · u**_*i*_. The resultant responses then compose a 50-cell set chosen as schematized in (d). The cells in {**u**_*i*_} are shown in (c) and their responses {*r_i_*} in (e).

**Figure 6:**
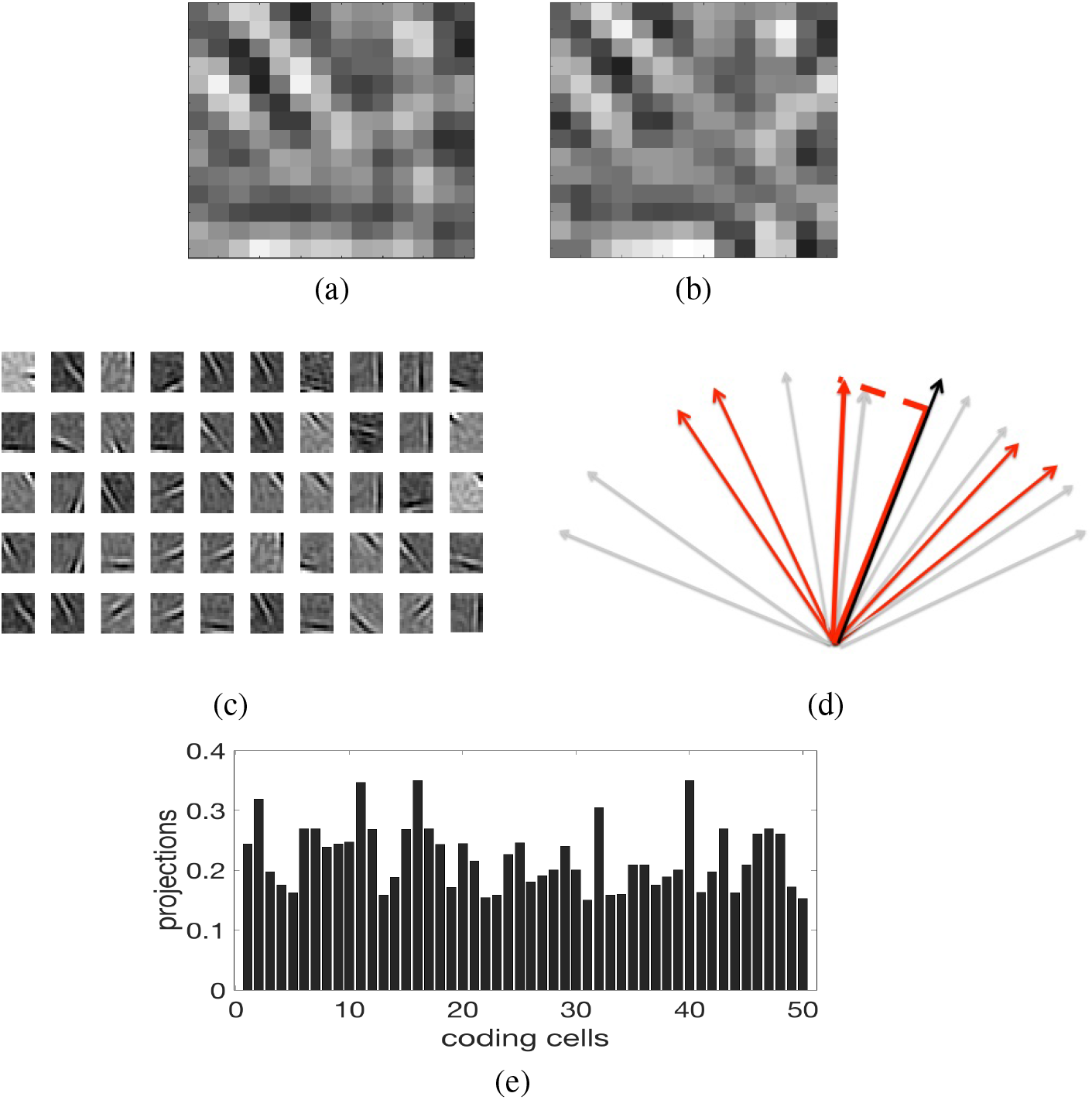
Parallel sparse coding model. (a) 196 filtered Lateral Geniculate Nucleus cells (LGN) provide input for a 14 × 14 pixel patch sampled from a natural image database. (b) The cortical reconstruction by summing 50 cells multiplied by their responses (c) The 50 cells that that were chosen by randomly sampling 1000 cells’ probability density function of the input projections. (d) Cells depicted abstractly as vectors are shown as gray. Ones that are chosen are shown in red and their length is their projection onto the input vector. Adding the red vectors and normalizing the results in a very good approximation of the input. (e) The 50 coding cells’ projections show their different magnitudes. The histogram of many such input codings is compared to experimental data. The similarities in the projections is characteristic of natural images.

Given that a single candidate neuron is chosen probabilistically, there are some details in processing the set of neurons that represent a stimulus within a single gamma cycle. Two implementation details are important to note. Firstly, as Figure 6c implicitly points out, the approximation is created by adding the cell responses and normalizing the sum [Carandini and Heeger, 2011]. We assume this can be done in parallel in the network. Secondly, there is the issue of choosing the cardinality of the cells. A reasonable approximation can be obtained from a binomial approximation of the distribution. In that case, the cells would be chosen from a pool of *N* neurons, where *r_ave_N* = 50, where *r_ave_* is the average response over a large population.

To make the final model histogram, one hundred images of LGN image patches are each coded with 50 projections and combined into a composite histogram, as shown in Figure 1 (b).

## 3 Discussion

Ever since the pioneering functional description of the action potential by Hodgkin and Huxley [Hodgkin and Huxley, 1952] models of action potential generation have become increasingly elaborate, e.g., [Prescott et al., 2008, Waters et al., 2005], and can account for increasing nuances of function as chronicled in standard texts, e.g., [Dayan and Abbott, 2001]. Nonetheless, there remains a significant gap between a cell’s action potential per se and its functional application in a behavioral context. In traditional analyses, some spatial or temporal population needs to be invoked to interpret the spikes’ functionality abstractly.

Our study has investigated how to extract delays in the action potential using a specific local gamma frequency reference. It provides concrete experimental evidence for Hopfield’s proposal and substantially extends [Vinck and Bosman, 2016, Vinck et al., 2010a], by introducing a coordinate system for the gamma cycle and associating the resultant delays with a sparse coding model. The distribution of phase delays of cells coding oriented grating images is closely matched to a sparse coding model encoding similar images. Essentially no translation is needed between delays and model responses since their messages form are in a scalar form used in standard neural models (as well as machine’s deep learning units). Much additional work needs to be done, but there is a possibility that this could be a standard way the cortex codes information.

One possible caveat is that the amplitude of the gamma frequency signal putatively controlling the action potential is very small. But one should keep in mind that the threshold governing the choice of the spike is also just a few milliseconds [Bean, 2007, Perrenoud et al., 2016]

Such a coding strategy has several potential advantages for a network over population-based models. One is the possibility to perform fast computations since the processing cycle can use the gamma frequencies directly. Additionally, as a delay code, it seems superficially compatible with spike-timing-dependent learning plasticity [Bi and Poo, 1998],but many potential propagation methods need to be examined e.g. [Lagache et al., 2019, Perrenoud et al., 2016, Stuart et al., 1997]. It is also compatible with multiprocessing [Ballard and Jehee, 2011, Ballard and Zhang, 2018], which reveals in a simulation that multiple gamma ‘clocks’ can coexist. Additionally, the compact coding of phase delays is compatible with sparse coding algorithms [Olshausen and Field, 1996a], which contain ready learning algorithms [Romo et al., 2004] including an extension to predictive coding [Rao and Ballard, 1999].

In 1995 it was generally accepted that the cortex signaled with populations of base one Poisson spikes [Sejnowski, 1995]. Although average firing remains extraordinarily useful, the phase code hypothesis argues that phase is more appropriate for a generative code and is not just a correlate. In comparison, population codes, despite their enormous utility in interpreting experiments, can face challenges as generative modes. In the mouse data, during the 3-second image presentation interval, the spiking rate is 0.2 spikes/second. In theory, this rate could be useful with population coding, but even moderate coding accuracy levels may need thousands of cells for modestly sized images, as shown in the supplementary material. Signals at gamma frequencies could be 100 ~ 1000 times faster than the current population coding strategies.

It is important to emphasize that allowing each spike to represent a separate analog quantity [Vinck and Bosman, 2016] is compatible with using the oscillatory frequency to group cells in a specific neuronal ensemble network [Vinck et al., 2010b]. To elaborate, a spike is either treated as a medium or a message in different contexts. The medium is the mechanism that synchronizes the process that is active on its neural network; the message is a quantity used by the recipient cell for the computation it is carrying out. [Panzeri et al., 1999] lists several message possibilities, such as a spike interval or a spike rate, but most of them are impractical owing to their computational inefficiency. Population coding is the least likely choice (but has its proponents [Akam and Kullmann, 2014]). Most schemes exclusively use the spike as a synchronizing mechanism, eschewing its ancillary use as a detailed message.

Finally, being able to associate computations with specific gamma frequencies may have even more advantages. Kobak et al. [Kobak et al., 2016] develop a variant of principal components analysis (dPCA) that forces the spikes that account for an experiment’s parameter variations to project on a small subset of axes. Data from Romo et al. [Romo et al., 2004] needed only 14% of the spikes to account for the task variance in their experiment. If it turned out that these spikes had a distinct gamma frequency [Fries, 2015], it would be one way that they could be distinguishable from the spikes in the much larger remainder.

The ability to pinpoint the role of specific cells in sub-populations could resolve many other issues in experiments. We can use the spikes in our data set as one example. The firing rates are very low, being 0.3 to 4 spikes per second, handicapping the network function’s explanation. Moreover, spikes measured outside of the stimulus ON interval have no obvious explanation in classical feed-forward terms but could have a potential explanation as an active two-way network embedded in the experimental context. Such issues are explored in a recent paper [Zhang and Ballard, 2020].

## Supplementary material

### Rate coding issues

Despite the advantages of volley phase coding, it is still common for rate averaging to be offered as a practical generative model [Panzeri et al., 2010, Akam and Kullmann, 2014]. However, there can be a significant cost with averaging codes. One might think that any precision needed could be achieved by having many cells use a small temporal window. However, given Poisson spikes, it takes a surprising number of cells to achieve modest levels of precision. To demonstrate this, we use the basic issue of estimating a scalar reliably from spikes that are generated with a probabilistic model. The model approximates a Poisson model but is elegant and shows that estimating a scalar accurately by counting spikes that are generated probabilistically takes a lot of them. The model has two parameters; 1) the desired accuracy and the probability of guaranteeing this level. Let’s use Hoeffding’s inequality [Hoeffding, 1994] where *m* spikes are modelled as *Z*_1_*, …, Z_m_* from a Bernoulli distribution where *P*(*Z_i_* = 1) = *ϕ* and *P*(*Z_i_* = 0) = 1 *− ϕ*. The estimate of the mean is given by 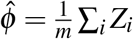. Let any *μ≥* 0 be a fixed error criterion. Then applying the inequality,

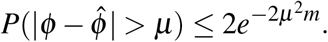

This formula allows one to estimate the number of cells to be counted to bound the error of the estimate. The rate/population model may need thousands of neurons or spikes to signal a scalar value reliably (within a reasonable error bound). There might be clever ways to save cells, but any direct way of producing an accurate numerical scale is going to be expensive.

### Additional data

The entire data set consists of 5 additional cells with 3456 spikes total. The data and related programs are freely available by contacting the second author.

## Notes

### Competing Interest Statement

The authors have declared no competing interest.

